# Assessing Stress Tolerance of *SUBI* and *DRO1* Introgression lines Under Flooding and Drought Conditions at Different Growth Stages

**DOI:** 10.1101/2024.09.23.614591

**Authors:** Ibrahim Soe, Emmanuel Odama, Alex Tamu, Aquilino Lado Legge Wani, Taiichiro Ookawa, Abdelbagi M Ismail, Jun-Ichi Sakagami

## Abstract

Rice varieties tolerant to submergence and drought regulate shoot elongation during short-term submergence by expressing the *SUB1A* gene, while the deep-rooted *DRO1* is effectively expressed under drought conditions to enhance water and nutrient uptake. This study investigates the growth and yield of rice with both *SUB1A* and *DRO1* in the background of IR64, under early season flooding and mid-season drought. The study used randomized complete design with two factors: soil moisture treatments (submergence, drought, and their combination) and genotypes. The genotypes included IR64, and three near-isogenic lines (NILs): NIL-SUB1DRO1, NIL-SUB1, and NIL-DRO1. Complete submergence was imposed for 7 days on 14-days old seedlings, while drought was imposed on control and submerged plants following a 21-day recovery period from submergence, using 42-day old plants. Variables were measured before and after treatments, and at harvest. The stresses negatively affected the genotypes. At harvest, IR64 and NIL-SUB1DRO1 under both stresses showed significant reduction in tiller numbers, shoot dry weights, and yields than their control plants. IR64 exhibited a significant delay in reaching flowering under all stresses. The rice introgression lines showed significant improvements of tolerance to the stress. The study showed no negative consequences of combining drought and submergence tolerance in rice.

## Introduction

Drought and submergence are considered the predominant abiotic stresses impacting global rice production (Lesk *et al*., 2016), affecting approximately 40 million hectares of rice-growing areas at various crop growth stages and negatively influencing plant growth, development, and yield (Barnabás *et al*., 2008). Although typically expected to occur at different levels in a topographic sequence, submergence and drought can occur in the same area within a single growing season in rainfed rice ecosystems.

Typically cultivated under partially flooded conditions, rice is a semiaquatic species. However, flash flooding that completely submerges rice plants for extended periods can cause most rice cultivars to die within a few days (Bailey-Serres *et al*., 2010). Flooding severely restricts O_2_ and CO_2_ gas exchange between rice tissues and the atmosphere, inhibits aerobic respiration and photosynthesis, and accelerates the consumption of energy reserves, leading to stunted growth and death (Kato *et al.,* 2014). Some rice genotypes can overcome complete submergence through antithetical growth responses. For instance, the submergence-tolerant lowland rice cultivar FR13A limits its elongation growth, conserving carbohydrate reserves to support the development of new leaves after desubmergence (Fukao and Bailey-Serres, 2008). This quiescence response is regulated by the *SUB1A* gene, which dampens ethylene production and enhances the mRNA and protein accumulation of two negative regulators of gibberellic acid signaling, resulting in the suppression of the energy-consuming escape response during submergence (Fukao and Bailey-Serres, 2008). Regeneration ability after flooding is vital for plants to produce high yields. Prompt regeneration of growth following flooding is an important plant characteristic (Panda *et al.,* 2008). After flooding, tolerant varieties show higher survival and faster regeneration ability by producing fresh leaves more rapidly and maintaining shoot growth (Panda *et al.,* 2008). Recovery is also faster in tolerant varieties due to their efficient reactive oxygen species scavenging systems and reduced lipid peroxidation when exposed to air (Singh *et al*., 2014).

Water unavailability in root zones is an important factor that reduces growth and yield of drought susceptible rice genotypes (Zhu *et al*., 2020). Alterations in agronomic traits under drought conditions alter the final crop yield, including the growth parameters, such as plant height and biomass at harvest, and components of harvestable yield, such as the panicle number per unit area, grain number per panicle, 1000-grain weight, panicle length, and filled grain percentage/seed setting rate (Zhang *et al*., 2018). Drought tolerance is defined as the ability of plants to survive under a low tissue water content (Kumar *et al*., 2016). Tuberosa (2012) demonstrated that drought-tolerant plants showed adaptive mechanisms such as higher leaf water potential and better osmotic adjustment, along with protective mechanisms such as leaf rolling and stomatal closure. Drought triggers the production or mobilization of the phytohormone abscisic acid, which is well-known for inducing stomatal closure to conserve water. Apart from abscisic acid, other factors that accumulate during drought and affect stomatal function include plant hormones (auxins, ethylene, brassinosteroids, and cytokinins), microbial elicitors (salicylic acid, harpin, and chitosan), and polyamines. Leaf rolling, a common adaptive response to drought stress in plants, is caused by alterations in water potential within epidermal and bulliform cells. Leaf rolling slows down transpiration and enhances water retention. Among the various components of drought tolerance in rice, root architecture remains a promising trait for analyzing differences in rice genotypes in response to drought, although it has been less explored than the above-ground changes (Gowda *et al*., 2011). The four primary root traits related to drought tolerance are root length, volume, thickness, and root growth angle (Uga *et al*., 2011). Root growth length is a key component of root traits for drought tolerance, as it determines root depth. Deeper and profuse root systems help plants survive drought stress by extracting water from deeper soil layers. In upland rice, these root systems enhance drought tolerance by improving the plant’s water uptake (Price and Courtois, 1999). Three major quantitative trait loci (QTLs) associated with root growth angles have been reported in rice (Uga *et al*., 2011). Among these three, DEEPER ROOTING 1 (*DRO1*) remains the major. Plant recoverability post-drought stress is also considered important, with some researchers suggesting its priority over drought tolerance itself (Sarkar and De Datta, 1975).

Despite the importance of submergence and drought to rice production in rainfed environments, the growth and yield of rice genotypes when subjected to both submergence and drought within the same season have not been fully investigated. Therefore, we evaluated the growth and yield of NIL-SUB1DRO1 developed in IR64 genetic background in the new rice lineage, under early flood and mid-growth drought. NIL-SUB1DRO1 was obtained by crossing NIL-SUB1 (a cross between IR64 and the submergence-tolerant rice FR13A) with NIL-DRO1 (a cross between IR64 and the drought-resistant rice Kinandang Patong). NIL-SUB1DRO1 was confirmed to contain both *SUB1* and *DRO1*.

## Materials and methods

This study was conducted from May to October 2023 in a greenhouse, with mean temperature and relative humidity of 27.9°C and 74.3%, respectively, during the experimental period.

The experiment comprised four environmental factors: control (optimum moisture), submergence, drought, and submergence–drought combination treatments. NIL-SUB1DRO1, NIL-SUB1, NIL-DRO1, and IR64 were the plant materials used in this study. The seeds were placed in a petri dish containing filter paper moistened with distilled water and left to germinate at 30°C in an incubator under dark conditions for 48 h. After germinating, the seeds were carefully selected and directly sown into polyvinyl chloride (PVC) pipes with an inner diameter of 8 cm and height of 80 cm, filled with 5.6 kg soil and arranged in a completely randomized design with 5 replications. During the initial submergence treatment, the top 20 cm of the 80 cm PVC pipe was cut off, and only this upper section was fully submerged. After the submergence treatment, the lower 60 cm section was reattached. To prevent nutrients deficiencies in the experimental soil, 10, 2.5, and 7.5 g of N, P, and K (8-8-8) compound fertilizer was mixed into the soil. For each treatment, three germinated seeds were sown per pipe, which were thinned to one seedling after five days.

Fourteen-day-old seedlings grown in 20 cm-high PVC pipes were transferred to a transparent glass container (80 × 55 × 50 cm) and completely submerged with tap water for seven days. Drought treatment was imposed on plants that received both submergence and drought treatments after 21 days of recovery from submergence. For plants that received only drought treatment, the treatment began after 41 days of seedling growth. Drought was induced by withholding irrigation for 14 days, while irrigation continued for the control plants and those that had previously undergone submergence treatment. At the panicle initiation stage, 0.4 g of nitrogen and 4.6 g of ammonium sulfate fertilizer were added to each PVC pipe.

## Measurement of variables

The greenhouse temperature and humidity were measured using the Long-Range Wireless Connection Logger Telemoni TML2101-A (AS ONE Corporation, Japan). A soil moisture sensor placed at a depth of 10 cm was used to measure the soil water content of the pipes. Data were recorded using a ZL6 Basic Datalogger (Meter Group, Inc. Pullman, WA, USA) with a 60-minute interval between each measurement throughout the drought period. Shoot length was measured from the base of the stem to the highest shoot tip using a meter ruler, taken before and after each treatment and at harvest. A chlorophyll meter (SPAD-502, Konica Minolta Corporation, Japan) was used to measure the chlorophyll content present in leaves. The SPAD value was an average of three measurements taken on the upper part of a newly fully developed leaf. This parameter was taken before and after each treatment, and at harvest. Days to flowering was determined by counting the days from sowing to when half of the culms of the plant in the PVC pipe had flowered (heading). Tiller and panicle numbers were determined by physically counting culm numbers and culms bearing grain, respectively. Tiller numbers were counted before and after each treatment and at harvest, while panicle numbers were only taken at harvest. To determine grain yield, the plants in each PVC pipe were harvested and threshed. After empty spikelets were removed, the filled spikelets were weighed. Following harvest, fresh shoots were separated from dry shoots and then oven-dried at 80°C for 72 h to determine the shoot dry weight. The root dry weight was determined by washing the roots with tap water, followed by oven-drying at 80°C for 72 h and weighing.

## Statistical analysis

Data were analyzed using a two-way analysis of variance. If significant differences were found, the least significant difference (LSD) test was conducted at *p* = 0.05 for mean separation.

## Results

Figure 1 presents the soil moisture content of the rice genotypes during the 14-day drought period. IR64, NIL-SUB1DRO1, and NIL-DRO1 showed reductions of 70.2%, 69.2%, and 70.5% in soil moisture content relative to the control respectively.

**Fig 1.**
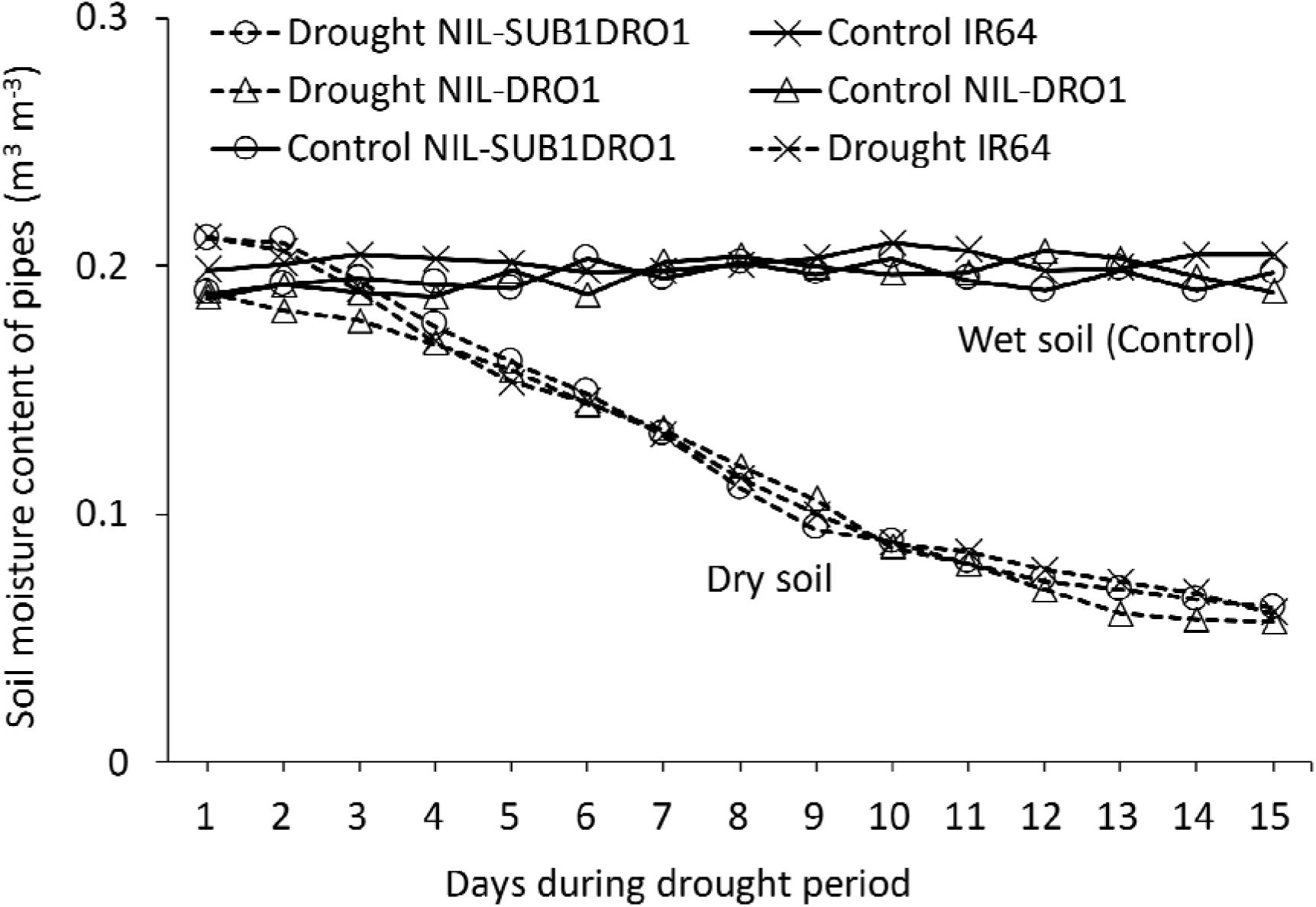
Soil moisture content of rice genotypes during the drought period.

The plant height of IR64 increased significantly by 22.1% during submergence, while those of submerged NIL-SUB1, NIL-SUB1DRO1, and control plants showed no significant difference (Figure 2a). At harvest, submerged IR64 showed a 17.7% decrease in plant height compared to the control. Under drought conditions, IR64 was significantly shorter than the control after 14 days of drought, while drought-treated NIL-DRO1 and NIL-SUB1DRO1 showed no significant difference in plant height compared to the control plants. At harvest, drought-treated IR64 and NIL-SUB1DRO1 showed respective 17.1% and 9.2% decreases in plant height compared to the control plants (Figure 2b). Under the combined submergence and drought treatments, the plant length of IR64 increased significantly during submergence but was markedly reduced pre-drought, post-drought, and at harvest compared to the control. IR64 showed a 12.9% decrease relative to NIL-SUB1DRO1 at harvest under a combination of the two stresses (Figure 2c). IR64, NIL-SUB1DRO1, and NIL-SUB1 showed respective decreases of 27.2%, 15.5%, and 16.5% in SPAD values after 7 days of complete submergence. At 14 days post-drought, no significant difference in SPAD values was observed between NIL-SUB1DRO1, NIL-DRO1, and the control plants, while IR64 showed a 14.6% decrease compared to the control plants (Figure 3b). Under the combined treatment, the SPAD value of IR64 decreased significantly, by 27.3% after seven days of complete submergence and by 8.6% after 14 days of drought. Similarly, NIL-SUB1DRO1 showed significant decreases of 15.5% after submergence and 5.6% after drought compared to the control. No significant differences were found between the genotypes and control plants for SPAD values under the different treatments during harvest.

**Fig 2.**
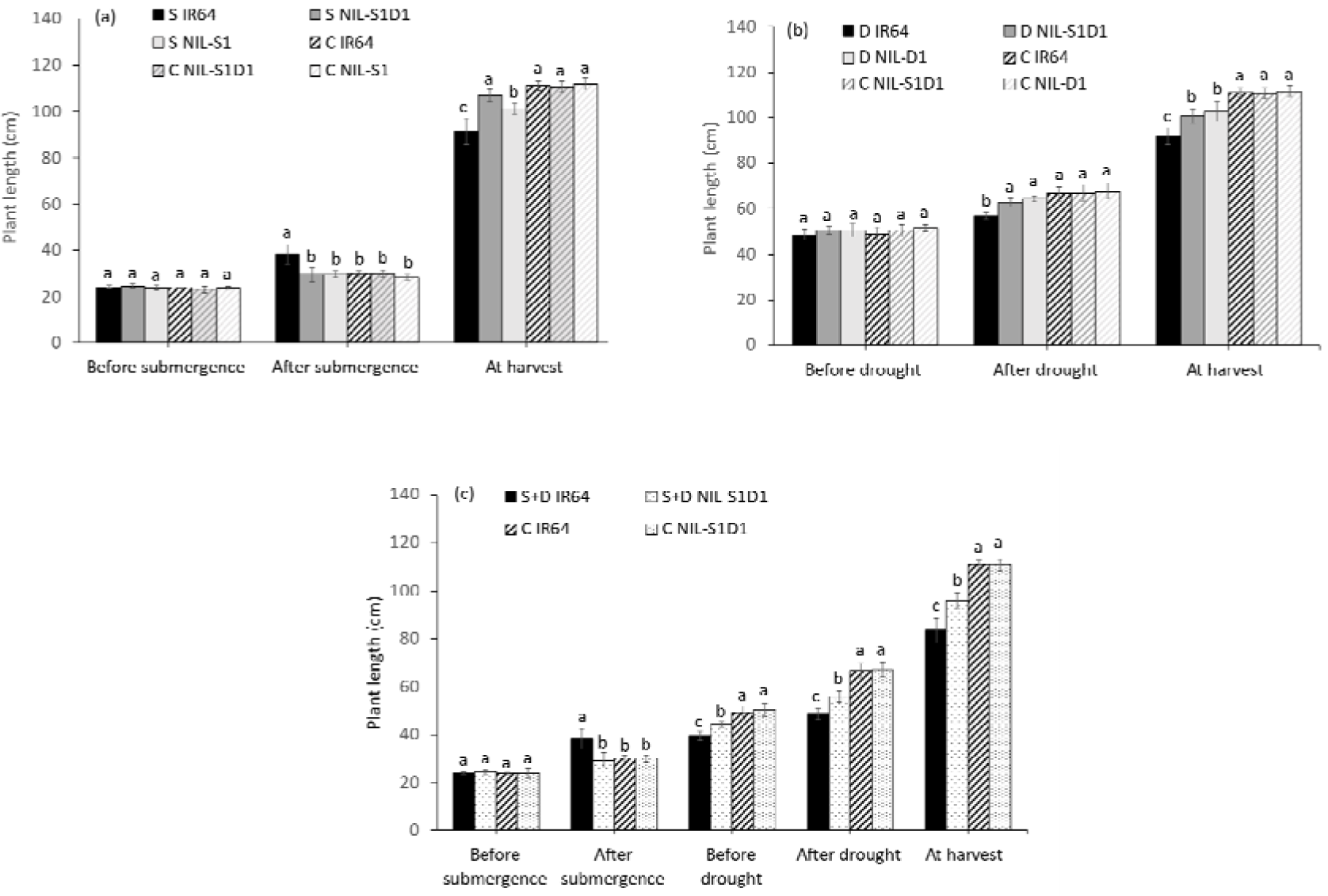
Plant length of submergence (a), drought (b), and submergence + drought (c) treatments. C = control, S = submergence, D = drought, S+D = submergence + drought, NIL-S1 = NIL-SUB1, NIL-D1 = NIL-DRO1, and NIL-S1D1 = NIL-SUB1DRO1. Different letters at the same time of data collection indicate a significant difference at *p <* 0.05 based on the least significant difference (LSD) test.

**Fig 3.**
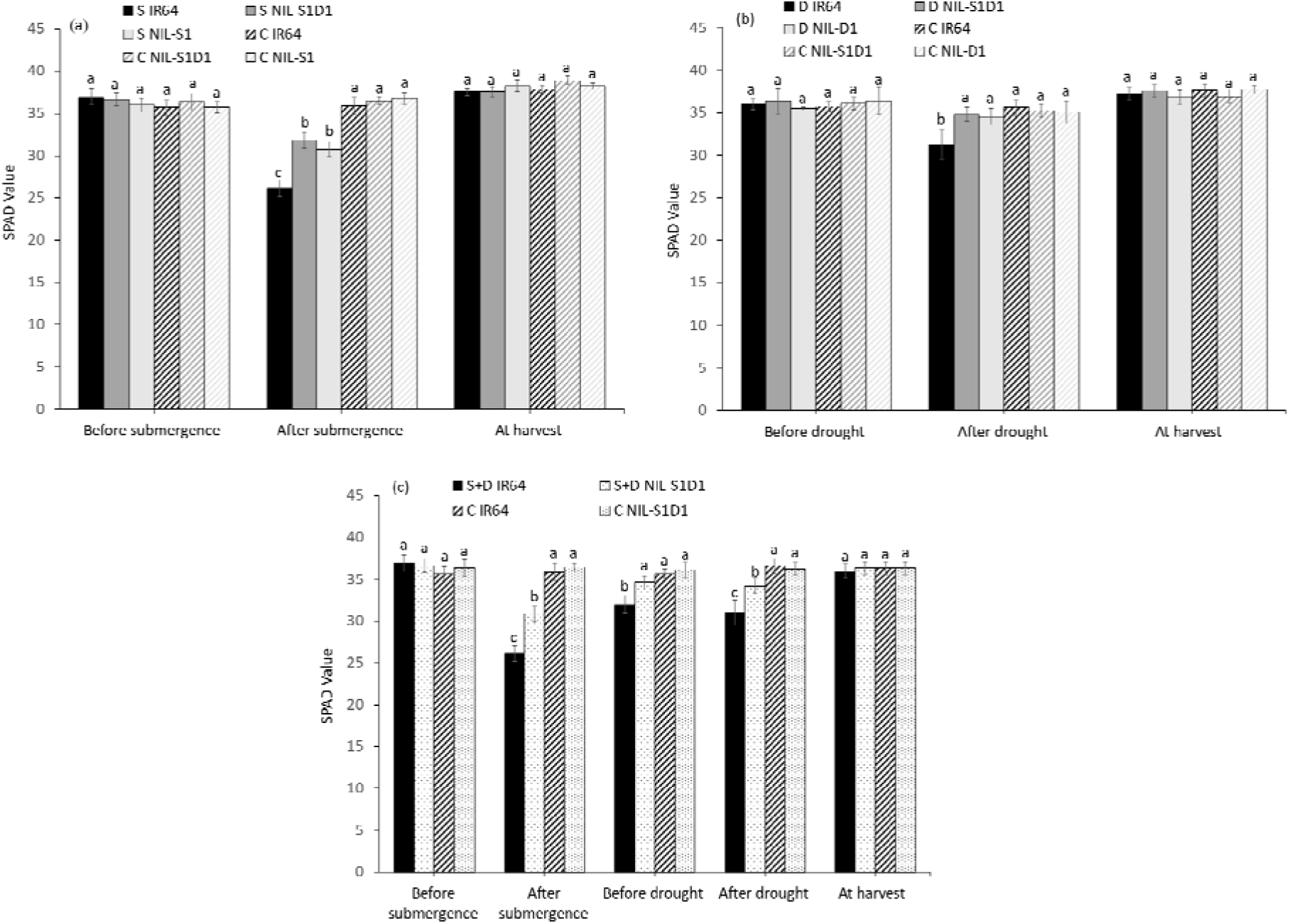
SPAD values of submergence (a), drought (b), and submergence + drought (c) treatments. C = control, S = submergence, D = drought, S+D = submergence + drought, NIL-S1 = NIL-SUB1, NIL-D1 = NIL-DRO1 and NIL-S1D1 = NIL-SUB1DRO1. Different letters at the same time of data collection indicate a significant difference at *p <* 0.05 based on the LSD test.

Before each of the three treatments, no significant differences were observed between the NILs in tiller number. However, IR64 showed 38.9% and 55.6% decreases under drought and combined treatment relative to the control. At harvest, submerged IR64 and NIL-SUB1DRO1 showed significant decreases in tiller numbers of 77.0% and 42.1%, respectively, relative to the control, with respective values of 66.3% and 37.5% under drought treatment and 71.9% and 51.1% under the combined treatment (Figure 4a, b and c). The different stresses delayed flowering of the genotypes compared to their controls. Submergence, drought, and their combination delayed flowering of IR64 significantly, by 10.4, 11, and 19.2 days, respectively (Figure 5). Significant differences were also observed between NIL-SUB1DRO1 and IR64 for days to flowering under submergence and drought, with IR64 showing significantly more days in both cases. No significant difference was found between the genotypes for days to flowering under control conditions.

**Fig 4.**
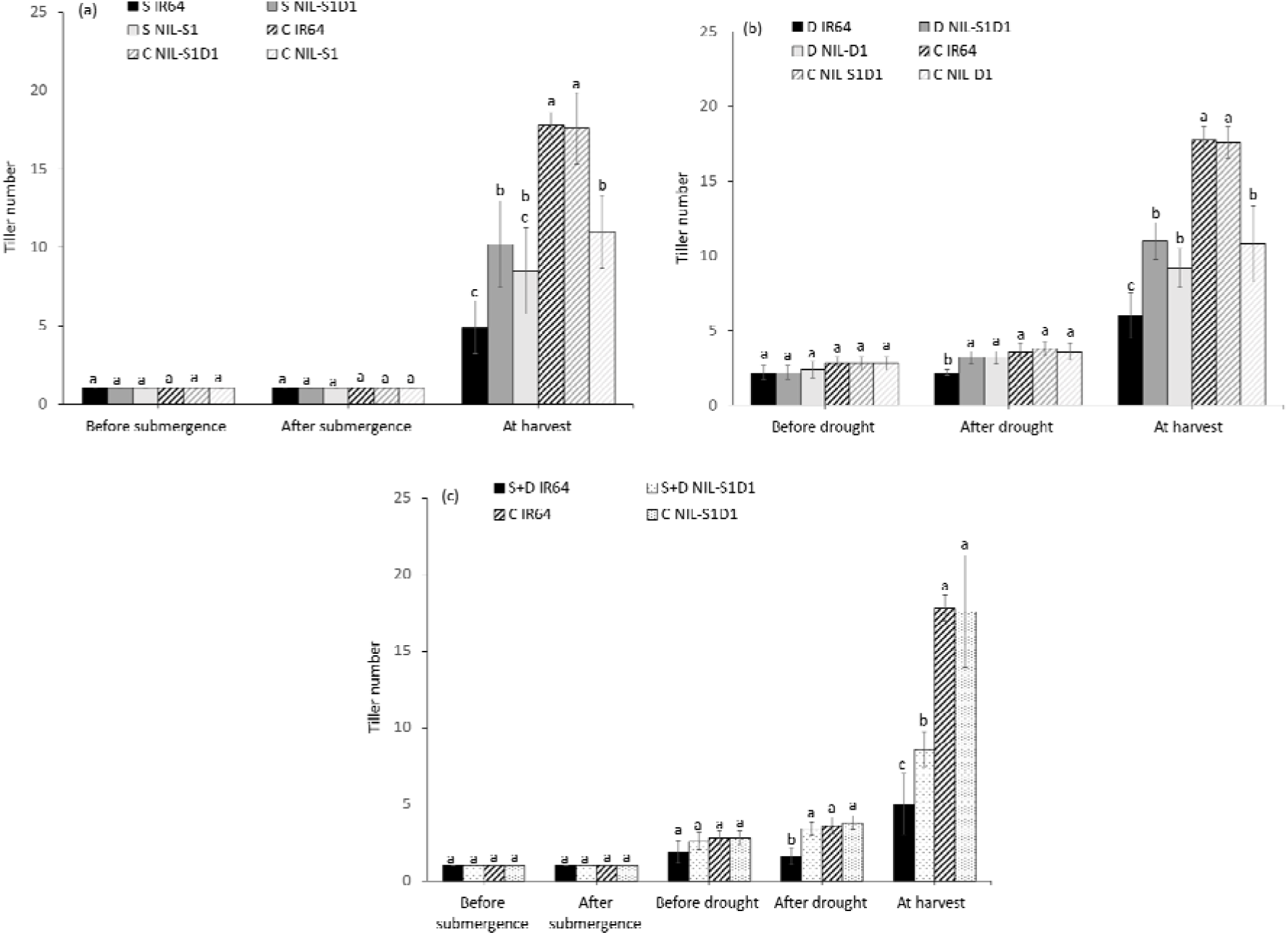
Tiller number of submergence (a), drought (b), and submergence + drought (c) experiments. C = control, S = submergence, D = drought, S+D = submergence + drought, NIL-S1 = NIL-SUB1, NIL-D1 = NIL-DRO1 and NIL-S1D1 = NIL-SUB1DRO1. Different letters at the same time of data collection indicate a significant difference at *p <* 0.05 based on the LSD test.

**Fig 5.**
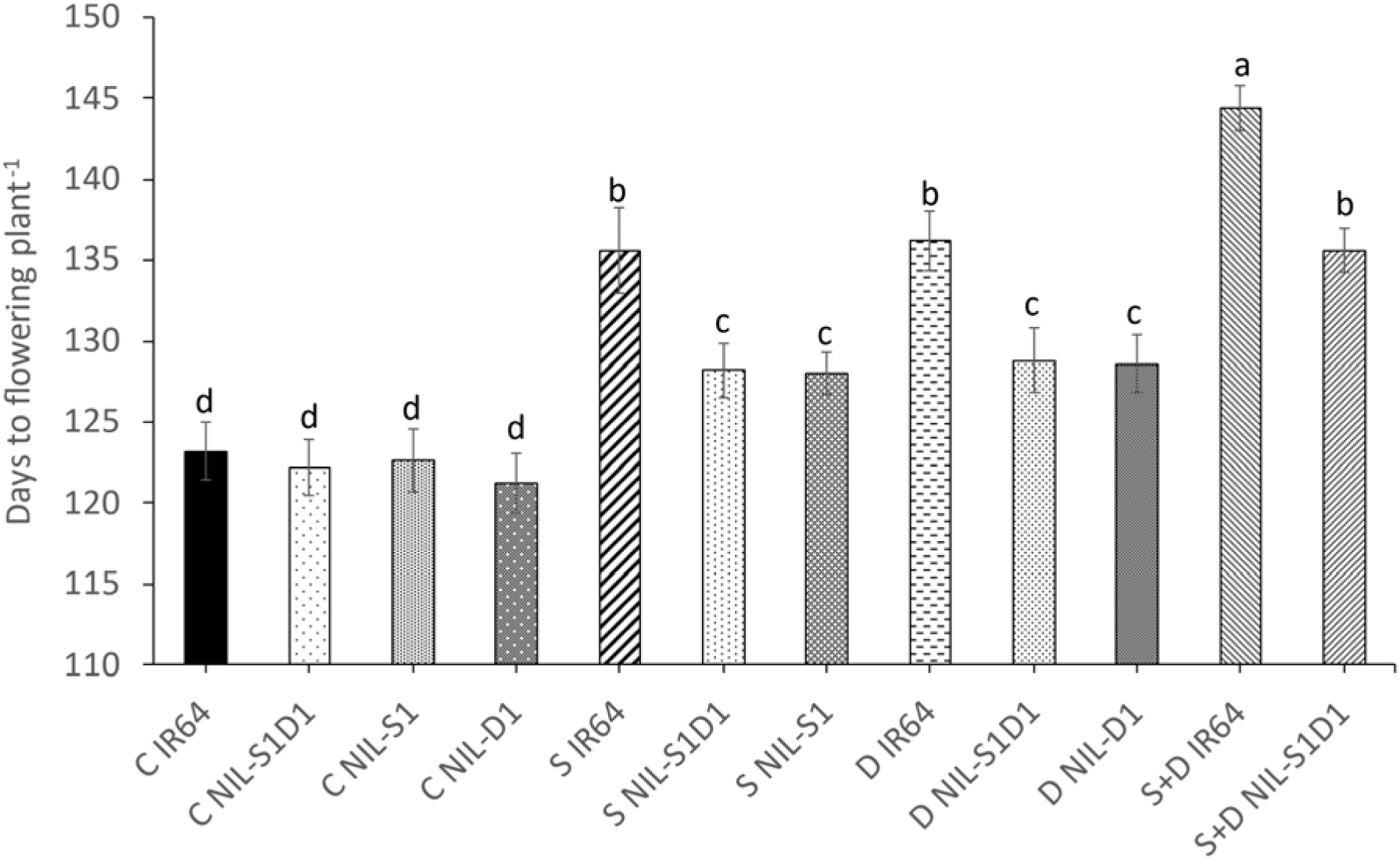
Days to flowering of submergence, drought, and submergence + drought experiments. C = control, S = submergence, D = drought, S+D = submergence + drought, NIL-S1 = NIL-SUB1, NIL-D1 = NIL-DRO1 and NIL-S1D1 = NIL-SUB1DRO1. Different letters indicate a significant difference at *p <* 0.05 based on the LSD test.

The different treatments markedly reduced yields of IR64, NIL-DRO1, and NIL-SUB1DRO1. Submergence, drought, and their combination reduced the grain yield of IR64 by 77.1%, 57.5%, and 75.5%, respectively, as well as that of NIL-SUB1DRO1 by 46.1%, 21.0%, and 52.8%, respectively, relative to the control (Figure 6). Drought significantly reduced the grain yield of NIL-DRO1 by 23.4%, and submergence markedly decreased the grain yield of NIL-SUB1 by 26.0%.

**Fig 6.**
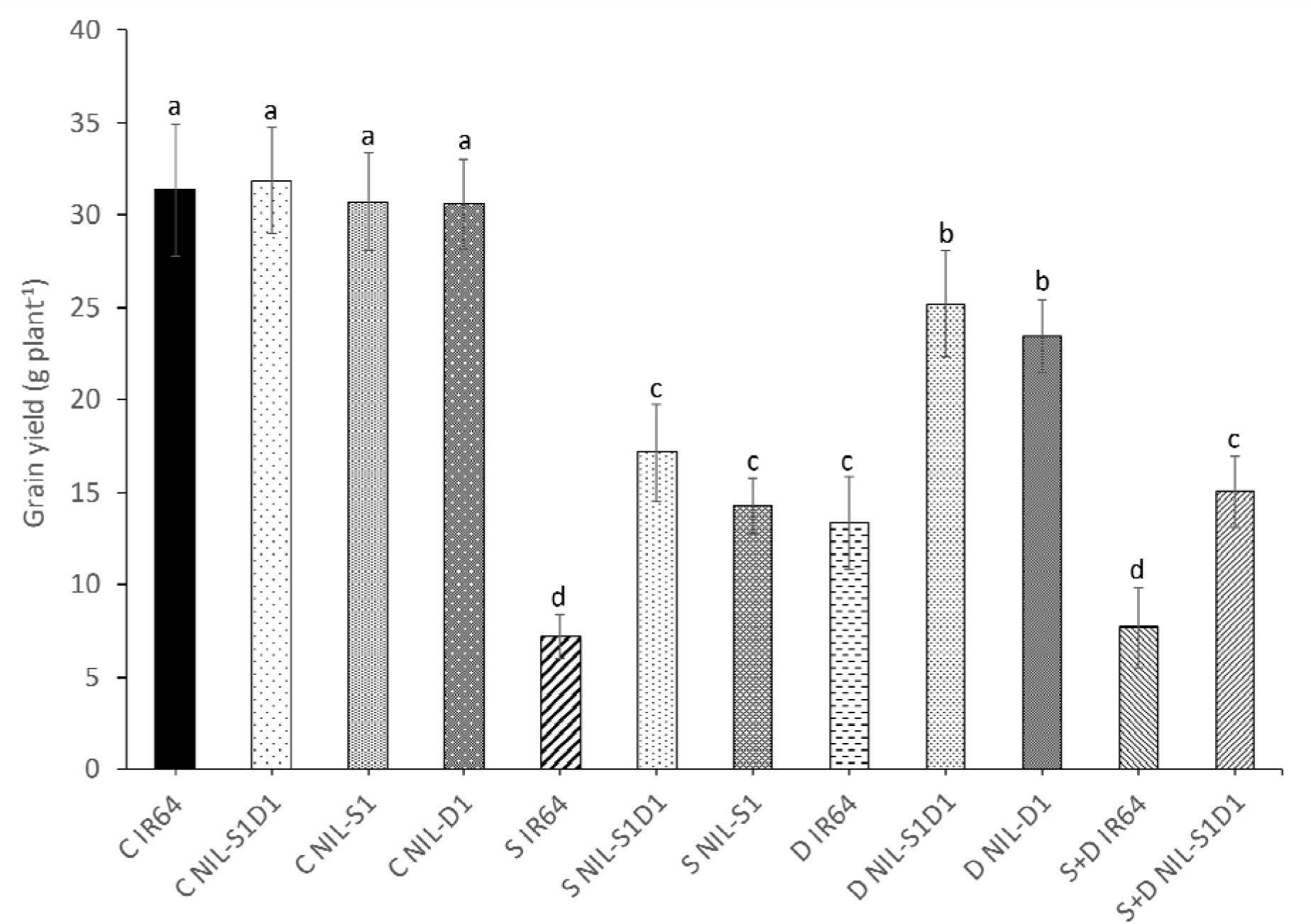
Grain yield of submergence, drought, and submergence + drought experiments. C = control, S = submergence, D = drought, S+D = submergence + drought, NIL-S1 = NIL-SUB1, NIL-D1 = NIL-DRO1 and NIL-S1D1 = NIL-SUB1DRO1. Different letters indicate a significant difference at *p <* 0.05 based on the LSD test.

Submergence significantly reduced the panicle number, shoot dry weight, and root dry weight of IR64 by 70.2%, 57.2%, and 65.2%, respectively (Tables 1 and 2). The panicle number, shoot dry weight, and root dry weight of NIL-SUB1DRO1 were also significantly reduced by submergence by 44.7%, 34.7%, and 32.7%, respectively. No significant difference was observed between the submerged and control plants of NIL-SUB1 for panicle number and shoot dry weight. Similarly, under drought treatment, no significant difference was found between the drought and control plants of NIL-DRO1 for shoot dry weight (Table 1). The panicle number, shoot dry weight, and root dry weight of IR64 were also significantly reduced under drought, by 63.8%, 40%, and 78%, respectively, while those of NIL-SUB1DRO1 were also reduced significantly but to a lesser extent, by 45.8%, 29.7%, and 40.0%, respectively. The combined treatment reduced the panicle number, shoot dry weight, and root dry weight of all genotypes, with respective reductions of 54.1%, 44.5%, and 27.3% for NIL-SUB1DRO1 and 72.5%, 63.7%, and 76.1% for IR64. The panicle length of IR64 was significantly reduced by 5.2, 20.6, and 8.3 cm following submergence, drought, and combined treatments, respectively. In contrast, no significant difference was observed in panicle length between drought-treated and combined stress-treated NIL-SUB1DRO1 and its control plants. Notably, the 1000-grain weight did not significantly differ due to submergence, drought or their combination, compared with the control plants.

**Table 1.** Shoot and root dry weights of IR64, NIL-SUB1, NIL-DRO1, and NIL-SUB1DRO1 after submergence, drought, and submergence + drought treatments.

| Treatment | Genotype | Shoot dry weight (g/plant) | Root dry weight (g) |
| --- | --- | --- | --- |
| Submergence | IR64 | 18.4 c | 3.2 c |
|  | NIL-SUB1 | 33.8 b | 6.8 b |
|  | NIL-SUB1DRO1 | 34.0 b | 7.4 b |
| Control | IR64 | 43.0 a | 9.2 a |
|  | NIL-SUB1 | 39.4 ab | 10.8 a |
|  | NIL-SUB1DRO1 | 45.8 a | 11.0 a |
| Drought | IR64 | 25.8 c | 2.2 c |
|  | NIL-DRO1 | 32.8 b | 6.6 b |
|  | NIL-SUB1DRO1 | 32.2 b | 7.2 b |
| Control | IR64 | 43.0 a | 10.0 a |
|  | NIL-DRO1 | 36.2 b | 10.8 a |
|  | NIL-SUB1DRO1 | 45.8 a | 12.0 a |
| Submergence + Drought | IR64 | 15.6 c | 2.0 c |
|  | NIL-SUB1DRO1 | 25.4 b | 9.6 b |
| Control | IR64 | 43.0 a | 9.4 b |
|  | NIL-SUB1DRO1 | 45.8 a | 13.2 a |

Different lowercase letters within a treatment indicate a significant difference at $p < 0.05$ based on the

**Table 2.** Yield components of IR64, NIL-SUB1, NIL-DRO1, and NIL-SUB1DRO1 after submergence, drought, and submergence + drought treatments.

| Treatment | Genotype | Panicle length (cm) | Panicle number | 100 grain weight (g/plant) |
| --- | --- | --- | --- | --- |
| Submergence | IR64 | 22.1 c | 4.2 c | 2.4 a |
|  | NIL-SUB1 | 28.3 a | 7.0 b | 2.2 a |
|  | NIL-SUB1DRO1 | 25.3 b | 9.4 b | 2.4 a |
| Control | IR64 | 27.3 a | 16.0 a | 2.4 a |
|  | NIL-SUB1 | 28.5 a | 8.2 b | 2.3 a |
|  | NIL-SUB1DRO1 | 28.6 a | 17.0 a | 2.4 a |
| Drought | IR64 | 22.0 b | 5.8 c | 2.4 a |
|  | NIL-DRO1 | 27.0 a | 9.2 b | 2.5 a |
|  | NIL-SUB1DRO1 | 27.3 a | 9.8 b | 2.4 a |
| Control | IR64 | 27.7 a | 16.0 a | 2.4 a |
|  | NIL-DRO1 | 28.5 a | 13.8 ab | 2.5 a |
|  | NIL-SUB1DRO1 | 28.7 a | 17.0 a | 2.4 a |
| Submergence + Drought | IR64 | 20.3 b | 4.4 c | 2.3 a |
|  | NIL-SUB1DRO1 | 25.0 a | 6.8 b | 2.5 a |
| Control | IR64 | 27.3 a | 16.0 a | 2.4 a |
|  | NIL-SUB1DRO1 | 28.6 a | 14.8 a | 2.6 a |

Different lowercase letters within treatment indicate a significant difference at $p < 0.05$ based on the LSD test.

## Discussion

The combined effects of submergence and drought are not considered to be greater than their individual effects. Therefore, this study aimed to analyze the growth and yield of rice with submergence and drought tolerance genes. NIL-SUB1DRO1 is a breeding line developed from crossing IR64, FR13A, and Kinandang Patong. IR64 is a modern lowland cultivar developed by the International Rice Research Institute and grown widely in South and Southeast Asia owing to its high yield and good grain quality, but it is sensitive to abiotic stresses, including submergence and drought. IR64 was primarily developed for irrigated rice production. An indica cultivar FR13A is highly tolerant of flooding and can survive for up to two weeks of complete submergence owing to a major QTL designated Submergence 1 (*Sub1*) near the centromere of chromosome 9. Kinandang Patong contains the deeper rooting 1 (*DRO1*), a major QTL involved in the deep rooting of rice under upland field conditions. This QTL controls the root system architecture and increases rice yield under drought conditions (Uga *et al*., 2011). Introducing rice varieties with submergence and drought tolerance to farmers is crucial, given the increasingly unfavorable climate changes.

Drought and submergence stresses are important abiotic factors that adversely affect the growth and yield of rice (Dixit *et al*., 2017). These stresses reduce plant growth by affecting various physiological and biochemical processes, such as photosynthesis, respiration, translocation, ion uptake, synthesis of carbohydrates, nutrient metabolism, and overall plant growth. In our study, submergence, drought, and their combination treatments markedly reduced the growth parameters and yield of the studied genotypes. Studies have shown that leaf photosynthetic rates of drought– and submergence-affected rice are reduced due to rapid declines in water availability, light intensity, and gas diffusion rates (Zhu *et al*., 2020). Consequently, the contents of non-structural carbohydrates in rice stems are reduced. Consistent with these studies, we found that submergence, drought, and their combination at the seedling and vegetative growth stages decreased the growth of rice genotypes, negatively affecting their grain yield. Grain yield is considered the most appropriate and direct criterion for selecting genotypes for drought tolerance due to the moderate to high heritability of this trait under drought conditions (Zain *et al*., 2014). Zhu *et al*. (2019) showed that the tiller number declined sharply during submergence, which supports our findings that tiller numbers of the genotypes were significantly decreased by submergence, drought, and their combination compared to the control. Accordingly, the lower tiller numbers led to reduced panicle number at the maturity stage.

A common adverse effect of water stress on crop plants is a reduction in fresh and dry biomass production. Farooq *et al*. (2009) showed that high dry weight under water stress conditions is a desirable characteristic for plant survival. This present study uncovered that the shoot and root dry weights of NIL-SUB1DRO1 were significantly higher than those of IR64 under stress conditions. Submerged IR64 showed significantly longer shoot after submergence but significantly shorter plants at harvest. According to the mechanism of submergence defined by Nagai *et al*. (2010) and Hattori *et al*. (2011), the elongation exhibited by IR64 represents transient submergence intolerance, whereas the relative quiescence exhibited by NIL-SUB1DRO1 represents tolerance. Shoot elongation during submergence, as experienced during flash flooding, is known to negatively impact flood tolerance due to the depletion of carbohydrates and the risk of lodging after desubmergence (Ram *et al*., 2002). NIL-SUB1DRO1 carries the *SUB1A* gene, which limits plant elongation during submergence, enabling a greater accumulation of shoot dry matter, compared to IR64 (Nurrahma *et al*., 2021). The significantly lower plant height of IR64 at harvest under the different stresses compared to control plants could be attributed to a decrease in turgor that impairs cell elongation and expansion (Ashfaq *et al*., 2022). Dessougi *et al*. (2022) reported that chlorophyll is the medium for absorption, transformation, and light energy transmission in plants; therefore, under abiotic stresses, plants with higher chlorophyll content or SPAD values have better chances of carbohydrate synthesis. A general delay in flowering occurs after submergence in all plant materials, as it takes the surviving plants additional time to recover and resume normal vegetative growth, and to repair the damage caused during and after submergence (Singh *et al*., 2009). In this study, the submergence-, drought-, and submergence–drought-tolerant genotypes, NIL-SUB1, NIL-DRO1, and NIL- SUB1DRO1, respectively, showed the least delay in flowering. This is likely because they experienced less damage from submergence, drought, and combined treatments compared to the sensitive variety, IR64. This shows that the introgression of *SUB1A* and *DRO1* narrowed the delay in flowering caused by submergence and drought by maintaining healthier plants during and after stress. The results also showed that combining *SUB1A* and *DRO1* does not have any negative consequences on rice growth and yield both under control and stress conditions, paving the way for breeding drought and submergence tolerant varieties adapted to rainfed lowlands.

## Conclusion

Submergence, drought, and their combination during early season flooding and mid-season drought significantly affected the growth and yield of the rice genotypes studied. However, NIL-SUB1DRO1, NIL-SUB1, and NIL-DRO1 maintained stable growth and yield. In particular, NIL-SUB1DRO1 was found to be resistant to the combined stresses of early flooding and mid-season drought during the growing season. The study findings provide empirical data that can inform breeding strategies to enhance rice submergence and drought tolerance in the future.

## Author contribution statement

Jun-Ichi Sakagami designed the study; Ibrahim Soe conducted the study; Taiichiro Ookawa provided rice genotypes NIL-DRO1 and NIL-SUB1DRO1, and Abdelbagi M Ismail NIL- SUB1; Emmanuel Odama, Aquilino Lado Legge and Alex Tamu assisted in data collection and analysis. Ibrahim Soe and Jun-Ichi Sakagami wrote the original draft. All authors participated in reviewing and editing of the manuscript.

## Acknowledgement

We thank Dr. Yusaku Uga of the National Agriculture and Food Research Organization, Japan for providing IR64DRO1 used to breed NIL-SUB1DRO1. The authors would also like to thank Enago (www.enago.jp) for the English language review.

## Abbreviations

LSD: Least significant difference
QTL: Quantitative trait loci

## Data availability statement

The article includes all data collected during this study. All the data in this study are available upon request.

